# The role of oxytocin in modulating self-other distinction in human brain: a pharmacological fMRI study

**DOI:** 10.1101/2021.10.17.463805

**Authors:** Yuanchen Wang, Ruien Wang, Haiyan Wu

## Abstract

The self-other distinction is crucial in human social cognition and social interaction. Studies have found that oxytocin (OT) sharpens the self-other perceptual boundary but with mixed results. Further, little is known if the effect of OT on self-resemblance facial perception exists, especially on its neural basis. Moreover, it is unclear if OT would influence the judgment in self-other discrimination when the other is a child or an adult. In the current double-blinded, placebo-controlled study, we investigated the effect of OT on self-face perception at both behavioral and neural levels. We morphed participants’ faces and strangers’ faces to create four stimuli conditions. After being treated by either OT or placebo (PL), participants reported whether a morphed face resembles themselves, or was morphed with their own faces, while being scanned with fMRI. Behavioral results showed that people judged adult-morphed faces better than child-morphed faces. fMRI results showed that the OT group exhibited generally increased activities in the visual area and IFG for self-morphed faces. Such difference was more pronounced in the adult face compared to child face conditions. Multivariate fMRI analysis revealed that the OT group showed better classification between self-morphed versus other-morphed faces, indicating that OT increased self-other distinction, especially for adult faces and in the left hemisphere. Our study shows the significant effect of OT on self-referential brain processes, providing evidence for the potential OT’s effect on a left hemisphere self network.

## Introduction

In human interactions, self-related and other-related information processing, such as self-other distinction and ingroup-outgroup separation, greatly contributes to everyday social decisions and interactions. When making these self-related or other-related judgments, one of the most important cue is the facial stimuli. Humans infer about genetic relatedness through the resemblance between the presented faces and self-face to adjust their altruism or investment in others’ behaviors (1, 2). Even though some studies have shown individual differences in processing self-resemblance faces (1–4), these faces also increased participants’ trust towards individuals by evoking the feeling of being close (5). Moreover, a study by Platek et al. also confirmed that implicit trust evaluation of self-resemblance faces would activate the reward-related brain areas (6), suggesting an important role of face in social interaction.

Successfully identifying and distinguishing information related to oneself and others plays a fundamental role in social life (7). Except for variance in stimuli (faces) themselves, administration of neuropeptide could also affect people’s related performance. For example, studies have shown that oxytocin (OT) promotes social connection and improves social interaction, such as trust and empathy (8, 9). However, how OT modulates the mentioned mechanisms and its impact on neural representation of social relationship is still unclear. One possibility is that OT decreases the self-other boundary and, in turn, increases social interaction. To further understand the mechanism of OT’s effect on social behaviors and fill the current research gap, it is crucial to investigate the effect of OT based on both behavioral and neural response during self-other face distinction tasks.

Given that OT can modulate the salience detection of social stimuli (10, 11), it is not surprising that OT influences face processing at both behavioral and neural levels (12, 13). For example, research showed that OT enhances recognition memory for faces but not for non-facial stimuli (14). Nevertheless, more recent evidence suggested that this enhancement is attributable to participants’ tendency to classify unfamiliar faces as familiar ones rather than an improvement in recognition memory for faces, as indexed by heightened sensitivity based on signal detection theory (SDT) (13). In addition, OT decreased amygdala responses to emotional faces, suggesting that OT tends to exert a greater impact on more socially salient information (15). Finally, among young male adults with autism spectrum disorder (ASD), OT was shown to increase participants’ tendency to fixate on socially salient facial features, such as the eyes (16). Taken together, these findings corroborate a role played by OT in increasing the social salience of facial features in face processing, which may in turn influences the representation of social relationship and further social behaviors.

Previous studies have shown that affiliative behaviors are supported by neural mechanisms associated with the social brain, including the medial prefrontal cortex (MPFC), the anterior cingulate cortex (ACC), the temporal parietal junction (TPJ), the inferior frontal gyrus (IFG), and the anterior insula (AI). (17–20). These brain areas are activated when individuals construct representations of relationships between the self and others and use this information to understand and guide social behavior (21–23). Therefore, it is possible that interpersonal psychological distance and self-other distinction are also mediated by the same brain network. Furthermore, the dorsomedial prefrontal cortex (dmPFC), one of the core regions of the social brain, is involved in mentalizing and social cognition (24, 25). dmPFC has been linked to social network size and the ability to create representations of the mental state of other individuals in both humans and primates studies (26, 27). Additionally, activity in this brain region has been linked to self-other distinction (26), and it is thought to depend on how close we perceive to other individuals and how similar we feel to them (28). Specifically, researchers have shown that dmPFC is activated when we make inferences about the mental state of dissimilar others as compared to similar others (29). Thus, dmPFC appears to be a core region that mediates social relationship and represents the mental state of other individuals, particularly when these individuals are dissimilar or unfamiliar. Similar to dmPFC, the ACC, the posterior cingulate cortex (PCC), and the TPJ have all been shown to play a part in mental state reasoning (21, 30). Previous studies showed that these regions can represent the position of the body in space and can help determine where an individual looks at (31). The ACC was also found to be active in self-monitoring behaviors, such as recognizing ourselves and others (31, 32). Together, these brain regions may serve as the neural underpinning of self-other distinction.

One’s own face is considered as salient self-related stimuli; therefore, it has long been applied in self-recognition and self-other perceptual differences studies (33, 34). Furthermore, from an evolutionary view, humans may show individual differences in detecting and expressing different feelings or behaviors regarding self-resemblance faces.The facial resemblance has been considered as a cue of human kin detection, which can be used to identify kinship relationships (2, 3). According to the inclusive fitness theory (35), humans would show increased prosocial tendency (such as investing, trustworthiness, or general attractiveness ratings) based on the closeness of kinship links. Researchers have shown that OT could modulate this self-resemblance face processing (36, 37), but with mixed results. For example, with self-stranger face morphing, a behavioral study indicated OT increased the ability to recognize differences between self and others in the self-other face differentiation task and increases positive evaluation of others (36). However, another result indicated that OT blurs the self-other distinction and reduces mPFC activity during self-trait judgments(37). Meanwhile, another important factor that influence the effect of OT is the age of the self-resemblance faces. Based on an fMRI study, child faces and adult faces were found to have different activation levels despite similar activation regions (38). Therefore, it is also crucial to consider the age factor in the current experiment and with faces of both adult and child involved.

Based on the current research finding in terms of the effect of OT on self-other distinction and the age factor, it could manifest two different hypotheses. Firstly, the OT effect on self-other distinction may be similar for both adult faces and child faces. Self-resembling faces indicate genetic relatedness and higher trustworthiness, which would activate reward-related brain regions, such as ventral superior frontal gyrus, right ventral IFG, and left medial frontal gyrus (MFG) (6). However, other evidence showed that OT increases self-other differentiation on peer-age faces (i.e., adult faces), but may not be evident for child faces. For instance, our previous study indicated that males are more sensitive to the self-morphed adult faces than self-morphed child faces (1). Thus, it is also possible that OT would increase self-other discrimination more under the adult-face conditions than the child-face conditions. The current study used a multi-modal pharmacological-fMRI approach to examine the neural correlates of the effects of OT on self-other distinction using self-morphed adult and child faces. In the study, we collected psychometric data on participants’ personality traits, behavioral data for the self-other discrimination task (e.g., accuracy and reaction time), and fMRI data of participants while executing the task. According to the social salience hypothesis of OT and own-age bias in face perception, we hypothesized that OT would affect self-other distinction and would be different for adult and child faces. In addition, we also expected that OT would modulate face-processing and self-processing brain activity depending on self-resemblance. Finally, we hypothesized that the OT effect on the self-other differentiation task (self vs. other) may be associated with one’s different personality traits.

## Methods

### Participants

We recruited 59 healthy male participants with the age range 20.9 ± 2.32 years, right-handed, and with 13 ∼18 years of education. They participated in this study via an online recruiting system. All participants filled out a screening form, and were included in the study only if they confirmed they were not suffering from any significant medical or psychiatric illness, not currently using medication, not consuming alcohol nor smoking on a daily basis. Participants were instructed to refrain from smoking or drinking (except for water) for 2 hours before the experiment. Participants received full debriefing on completion of the experiment. Written informed consent was obtained from each participant before experiment. The study was approved by the local ethics committee.

### Task design

In the study, we divided participants into two groups, with one of them treated with nasal oxytocin administration (OT group) and the other treated with placebo (PL group). For the stimuli presented to each participant, we created four experimental conditions (self-child, self-adult, other-child, and other-adult) by morphing the participant’s face with one of two adult faces with neutral expression (a 23 years old male face or a 23 years old female face, according to the participants’ gender) and a 1.5 years old child face. Fig. 1 shows the illustration of morphed faces and resulting four experimental conditions, a similar process as our previous study (1).

**Fig. 1.**
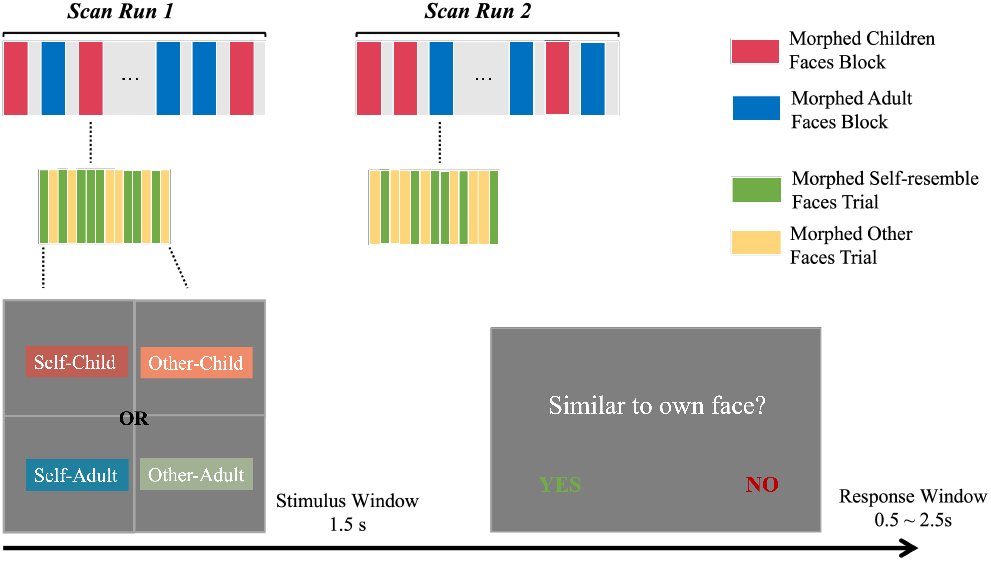
Sequence of the experiment. Different conditions are color coded indicated by legend in the upper right corner. Faces shown in the Self-Child condition were morphed using the participant’s own face and a stranger child’s face; faces in the Other-Child condition were morphed using an adult stranger’s face (the same gender as the participant) and a stranger child’s face; faces in the Self-Adult condition were morphed using the participant’s own face and a stranger adult’s face (the same gender as the participant); faces in the Other-Adult condition were morphed using two adult strangers’ faces (the same gender as the participant).

We collected data in two six-minute runs for each participant. In each run, there were six blocks, including three adult blocks and three child blocks that presented in a randomized sequence. In the adult blocks, the stimuli shown were those morphed with adults: the self-adult and other-adult conditions. Similarly, only self-child and other-child conditions were in the child blocks. For each trial in the study, the facial stimuli were presented for 1500ms, which shows the target morphed face. Participants were asked to judge if the face resembled with their own faces in the following response window (Fig. 1).

### Material preparation and acquisition

The full-face photograph of each participant was taken one week before the formal study. The photographs were taken 3 days before the scanning day. Participants were asked to keep neutral expression when facing the camera. The morphed faces were created based on those taken photographs.To exclude the gender effect of the child face, we did a gender rating task to the child face in a 5 point scale (1 = a girl, 2 = maybe a girl, 3 = not sure, 4 = maybe a boy, 5 = a boy), and the rating result indicated that both male (mean rating = 3.17, SD = 1.47) and female (mean rating = 2.69, SD = 1.13) participants showed uncertainty of the gender of the child face. Additionally, faces used in the other-adult condition and other-child condition were the same for female and male participants. All faces were processed with Adobe Photoshop CS to standardize the picture to black and white, with merely interior characteristics of face being retained. Then the Abrosoft Fanta Morph (www.fantamorph.com) software was used to create the 50/50 morph of the two selected faces, the similar method as used in previous studies (6, 39, 40). Thirty calibration locations were used to make the morphed face in a standard face space and all output morphed faces were resized to 300 × 300 dpi. All stimuli were presented on a 17-inch Dell monitor with a screen resolution of 1024× 768 pixels and 60 Hz refresh frequency, the visual angle of the face images is 4.3° ×4.6° and the mean luminance of stimulus was 166 *cd/m*^2^.

### Psychological scales

We collected demographic and psychometric data of all participants. The measures was predominantly recorded before the experiment, with several exceptions. “PA after” “NA after” and “SA after” are the three measures taken after OT/PL administration. These measures used the same scales as PA (positive affect), NA (negative affect), SA (social anxiety), which were taken before the treatment (Table 1). The psychometric data was collected as a composite of control factors to make sure that people administered OT have no significant difference from people administered PL. For example, Interpersonal Reactivity Index (IRI) is a widely-used assessment of empathy (41). There are four subscales in IRI, including perspective taking (PT), fantasy (FS), empathetic concern (EC), and personal distress (PD). Each subscale includes seven questions. EC measures individuals’ feelings of compassion and concern for others. FS describes the tendency that respondents transpose themselves into fictional characters. PD indicates the extent that individuals feel uneasiness when exposed to the negative experiences of others. PT assesses unplanned attempts to adopt others’ points of view. Mean scores were subsequently compared across treatment groups (OT, PL) to rule out effects of OT on these measures. Because of our randomized design, we predicted that there would be no significant difference between OT and PL group in psychometric scores.

**Table 1.** Psychometric data and questionnaire scores

|  | PL | OT | t-score | p-value |
| --- | --- | --- | --- | --- |
| PA | 31.76 (5.33) | 33.1 (6.99) | -0.8242 | 0.4136 |
| NA | 18.17 (6.12) | 17.17 (6.02) | 0.6272 | 0.5331 |
| SA | 37.24 (9.29) | 35.03 (8.88) | 0.9248 | 0.3591 |
| PA after | 29.38 (5.73) | 31.21 (6.86) | -1.1009 | 0.2758 |
| NA after | 17.48 (6.65) | 16.69 (6.2) | 0.4697 | 0.6404 |
| SA after | 38.69 (9.1) | 37.34 (10.73) | 0.5148 | 0.6088 |
| AQ | 22.14 (4.84) | 23.59 (5.64) | -1.0494 | 0.2986 |
| Neuroticism | 44.97 (12.01) | 40.31 (11.75) | 1.492 | 0.1413 |
| Extraversion | 48.9 (10.3) | 51.31 (10.42) | -0.8873 | 0.3787 |
| Openness | 55.86 (8.48) | 54.69 (7.97) | 0.5426 | 0.5896 |
| Agreeableness | 61.86 (6.58) | 59.24 (8) | 1.3619 | 0.1789 |
| Conscientiousness | 60.52 (11.66) | 63.52 (10.18) | -1.0439 | 0.3011 |
| IRI | 96.76 (9.64) | 92.17 (10.17) | 1.7619 | 0.0836 |
| TAS-20 | 53.55 (9.3) | 49.48 (9.54) | 1.6443 | 0.1057 |
| BIS all | 66.59 (8.83) | 65.59 (9.81) | 0.4079 | 0.6849 |
| SES score | 30.03 (4.36) | 31.62 (3.86) | -1.4666 | 0.1482 |
| NPI score | 121.24 (25.38) | 127.07 (25.25) | -0.8766 | 0.3845 |
| SDO score | 53.45 (12.45) | 55.72 (10.53) | -0.7517 | 0.4555 |
| FNE short | 42.28 (7.43) | 39.93 (7.45) | 1.2001 | 0.2351 |
| FNE all items | 103.86 (15.44) | 97.52 (15.84) | 1.545 | 0.128 |
| Mav score | -3.34 (11.53) | 1.76 (14.14) | -1.5064 | 0.1378 |
| father love | 23.34 (9.42) | 22.66 (7.58) | 0.3072 | 0.7598 |
| mother love | 23.03 (9.58) | 24.93 (7.64) | -0.8334 | 0.4083 |
| NFCC | 163.24 (16.36) | 160.93 (18.92) | 0.4974 | 0.6209 |
| IUS All | 74 (18.54) | 75.55 (16.48) | -0.3369 | 0.7374 |
| IUS Chinese | 66.83 (16.04) | 67.69 (14.14) | -0.2171 | 0.829 |
| Perspective taking | 22.48 (4.04) | 22.34 (2.32) | 0.1594 | 0.8741 |
| Fantasy Scale | 25.17 (5.52) | 23.14 (5.4) | 1.4188 | 0.1615 |
| Empathic concern | 26.76 (3.66) | 25.28 (4.84) | 1.3158 | 0.194 |
| Personal distress | 22.34 (3.92) | 21.41 (4.37) | 0.8538 | 0.3969 |

**Note:** Means(standard deviations), and two-tailed t-test (direction: OT - PL) results of the comparisons on demographics data. As non of the comparisons were significant, multiple comparison corrections were not applied.
PA: positive affect; NA: negative affect; SA:social anxiety; AQ: autism quotient; IRI:interpersonal reactivity index; TAS-20:Toronto Alexithymia Scale; BIS: behavioral inhibition system scales; SES: socioeconomic status; NPI:Narcissistic Personality Inventory; SDO: social dominance orientation; FNE: fear of negative evaluation; Mav score: Machiavellianism Scale; NFCC: need for closure scale; IUS: intolerance of uncertainty scale

### fMRI Image collection

All images were acquired on a 3T Siemens Tim Trio scanner with 12-channel head coil. Functional images employed a gradient-echo echo-planar imaging (EPI) sequence with following MRI scanner parameters:(TE = 40*ms*, TR = 2*s*, flip = 90°, FOV = 210*mm*, 128 by 128 matrix, 25 contiguous 5*mm* slices parallel to the hippocampus and interleaved). We also acquired the whole-brain T1-weighed anatomical reference images from all participants (TE = 2.15*ms*, TR = 1.9*s*, flip = 9°, FOV = 256*mm*, 176 sagittal slices, slice thickness = 1*mm*, perpendicular to the anterior-posterior commissure line).

### fMRI Imaging analysis

fMRI data preprocessing was performed using Statistical Parametric Mapping software (SPM12: Wellcome Trust Centre for Neuroimaging, London, UK). The functional image time series were preprocessed to compensate for slice-dependent time shifts, motion corrected, and linearly detrended, then coregistered to the anatomical image, spatial normalized to Montreal Neurological Institute (MNI) space (http://www.bic.mni.mcgill.ca/ServicesAtlases/HomePage) and spatially smoothed by convolution with an isotropic Gaussian kernel (FWHM = 6 mm). The fMRI data were high-pass filtered with a cutoff of 0.01 Hz. The white matter (WM) signal, cerebrospinal fluid (CSF) signal and global signal, as well as the 6-dimensional head motion realignment parameters, the realignment parameters squared, their derivatives, and the squared of the derivatives were regressed. The resulting residuals were then low-pass filtered with a cutoff of 0.1 Hz.

### Univariate analysis and ROI analysis

Univariate analysis was conducted using the SPM toolbox with general linear models (GLM) (42). In the GLM analysis, face stimuli blocks were modeled with a box-car function, convolved with a standard hemodynamic response function. Four conditions were defined by separate regressors: self −adult, other −adult, self −child, other −child. Six head movement parameters from the spatial realignment were entered as covariates of no interest. Statistical parametric maps were generated for each subject from linear contrasts between each of the four conditions. For the ROI analysis, ROIs were defined based on coordinates from prior literature indicating significant dynamic activity in self-resemblance face processing (43). The ROIs were selected and then labeled using the *xjV iew* toolbox (https://www.alivelearn.net/xjview). Moreover, masks from Neurosyth meta-analysis were also obtained for self-related masks using the keyword “self”. ROI masks from AAL3 (44) were constructed by resampling the AAL3 template using affine transformation. The percentage signal changes (PSC) over these ROIs were calculated based on the work of Mazaika 2009 (45). We tested our main hypothesis by calculating condition differences between PSC in the self face processing ROIs and social brain ROIs.

### Multivariate brain analysis

Previous literature showed that univariate and multivariate analysis on the fMRI would result in similar but not identical information on patterns of brain activation (46). Therefore, it is important to analyze the data using multivariate methods to support and strengthen any discoveries found in the univariate processes. It is also possible that more hidden detail about how OT influence brain activation would be revealed through the further analysis of multivariate methods.

### Principal component analysis (PCA)

Because of the high dimensionality of fMRI data, we have conducted PCA for dimensionality reduction and identifying OT effects in task-based fMRI. Specifically, we performed spatial PCA on task fMRI activities and extracted the first fMRI component that explained the most variance in data. We then investigated importance of psychometric scores to the projection of this principal component (Fig. 13). Principle component and projections were generated using *sklearn* package (47). Reported principle component (Fig. 13) were based on the activation pattern of all participants.

### Multivariate pattern analysis (MVPA)

Unlike traditional analysis using univariate or mass-univariate approaches, the MVPA considers patterns of responses across multiple voxels, rather than single voxel-based or region-based values. The fMRI data is naturally multivariate, which allows for the multivariate analysis of multi-dimensional data. MVPA is a machine learning based approach, mainly dealing with classification and regression problems. In this way, the activation of thousands of voxels in fMRI data is reduced to accuracies in several classifiers. In the current study, the analysis was conducted using the *Decoding* Toolbox (TDT) (48). We first applied the boxcar function from the preprocessed fMRI data producing four beta estimates per run for each participant. Each of the four beta estimates was then correlated to the self-child, self-adult, other-child, and other-adult conditions, respectively. After the correlation, we did a group-level MVPA from the generated beta images. Under each one of the four conditions, we performed a whole-brain MVPA on discriminating OT or PL treatment with the searchlight method. The goal was to generate the accuracy of each voxel on discriminating the treatment.

### Representational similarity analysis (RSA) and representational connectivity analysis

To understand the similarity and difference between two group in conditions, we used the *NeuroRA* toolbox (49) to extract representational dissimilarity matrices (RDM). We chose the same ROIs as used in the PSC analysis and calculated the RDM for each ROI from the OT or the PL group, respectively. Then, we were able to derive the correlation matrix among the ROIs. For comparison between the two correlation matrices, we used the Fisher-z transformation to derive the significance of the OT and PL ROI-connectivity difference (see Fig. 10).

## Results

### Psychometric Data

The questionnaire scores of participants in the two treatment groups were summarized in Table 1. The means and standard deviations of the scores were calculated among each group, separated by treatments (OT and PL). We also performed pairwise comparison statistics between OT and PL treatment of these questionnaire scores (Table 1). The full names of each score were listed in the table caption. None of the two-tailed *t*-tests were significant, providing no evidence for group differences between OT and PL treatment in participants’ psychometric data. Furthermore, participants were asked to take each of the PA, NA, and SA tests twice, with one before the treatment administration and one after the treatment administration. The absence of significance changes provides no evidence regarding the role of OT on these trait, or emotional state scores.

### Behavioral performance

Supposing the correct answer for self-morphed faces is “yes” (see Task Design for more details), we derived four conditions (namely, OT-child, OT-adult, PL-child, and PL-adult) and calculated the accuracy separately (Fig. 2(a)). We calculated overall accuracy for each participant and put into a two-way mixed ANOVA using treatment groups (OT vs. PL) as between-subject factor and facial conditions (self vs. other, and child vs. adult face) as within-subject factors. The result of ANOVA showed a significant main effect of facial conditions (*F* (1, 57) = 54.6716, *p*<0.001, 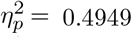), where participants showed higher accuracy on adult face discrimination (Fig. 2(a)). However, there is no significant effect of treatment (*F* (1, 57) = 0.1119, *p* = 0.7392, 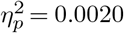). These results are consistent with previous literature, which indicate that the participants could better detect faces at their own age (1). No other significant effects on accuracy were identified, *p*-values *>* 0.05.

**Fig. 2.**
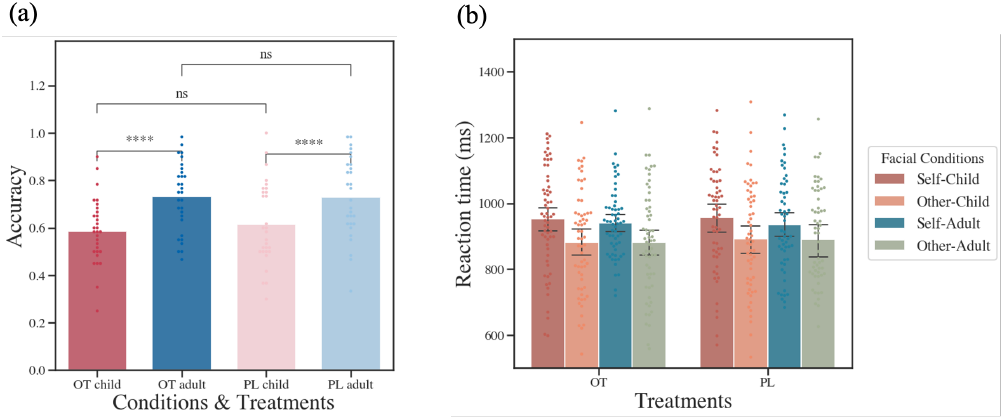
Accuracy and reaction times of each condition. (a) The mean accuracy under each condition for the two groups. The bars indicate the average accuracy for that condition, while the dots show accuracy of each participant in each run. The pair-wise comparisons are labeled in the figure, with “***” indicating 1 · 10^*−*4^ *< p <* 1 · 10^*−*3^, and “ns” indicating 0.05 *< p <* 0 (non-significant). Neither differences of the between-group comparisons (OT vs placebo) is significant, while both differences of the within group comparisons (child vs adult face) are significant. The result of these *t*-test corresponds to the F-tests in ANOVA. (b) The reaction times of participants, bars are divided by treatment and facial conditions of the stimuli.

For reaction times (RTs), similar analysis indicated that participants exhibited a significantly longer RTs to discriminate self-morphed faces than other-morphed face (OT group: *F* (1, 29) = 18.6276, *p* = 0.0002, 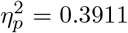, PL group: *F* (1, 28) = 10.3001, *p* = 0.0033, 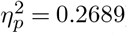). No significant effects on child/adult face on RTs was observed (Fig. 2 (b)). To test whether there was RT difference for responding as “self” or “other” in two groups, we also conducted ANOVA on RTs with age (adult vs. child) by response type (responded as self vs. respond as other). It did not show significant treatment effect, but suggested longer latency for responding as “self” for child faces than other types of faces (Figure S1).

As there were multiple conditions, overall accuracy would be a over-simplified generalization to the behavioral data. Confusion matrices for each participant was constructed, and we calculated the true positive rate and false positive rate from the confusion matrices. To better visualize the decisions of participants under each condition, we fitted the receiver operating characteristic (ROC) curve for each condition and calculated the detectability (d’) and criteria (c) for each participant based on SDT. The ROC can illustrate the diagnostic ability for a specific condition, where we then compared the curves and their area under the curve (AUC) to see if there is any significant difference. Pair-wise significance comparisons showed that higher AUC for adult conditions than child conditions, indicating better performance to detect their own faces in adult conditions (Fig. 3). The ROC curve results were consistent with accuracy results, showing no significant effect on other factors. Furthermore, based on SDT and the work of Dal Martello & Maloney (2006) (50), we computed the detectability and response criteria for each participant. However, as shown in the results of Fig. 4, none of the comparisons is significant.

**Fig. 3.**
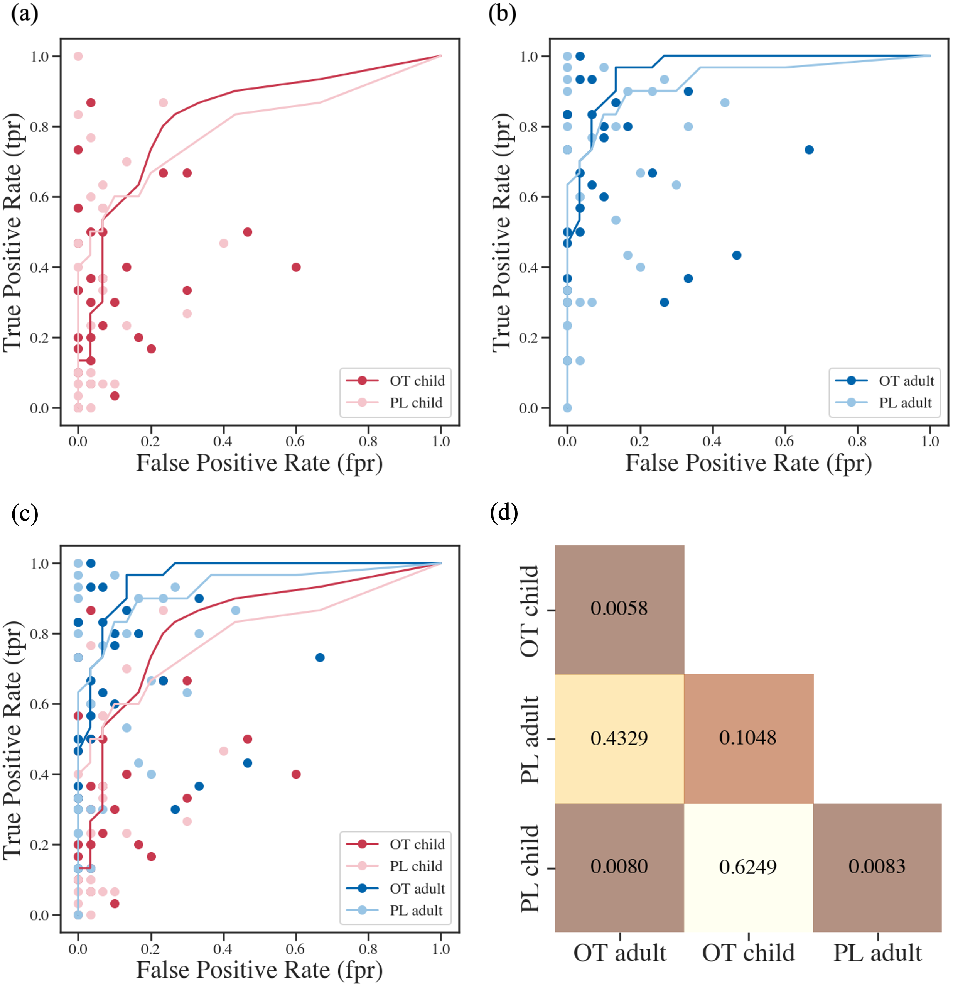
Fitted ROC curves and *p*-values of the pair-wise comparisons. (a) ROC curves for the child face conditions. (b) ROC curves for the adult face conditions. (c) Composite ROC plot for all four conditions. (d) Heat-map of *p*-values for pairwise ROC comparisons. *p*-values are labelled in the box for each corresponding comparisons. As indicated in the heat-map, the only significant comparisons are: OT child OT adult, Placebo child Placebo adult, and PL child OT adult. This result is consistent with the ANOVA and *t*-tests results, such that the behavioral results indicated significant adult vs. child difference, but no treatment effect.

**Fig. 4.**
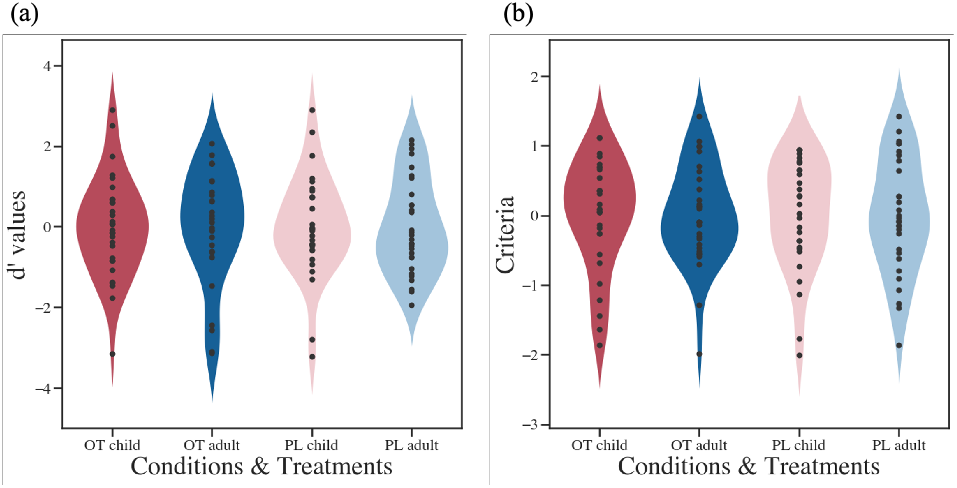
(a) The *d*^*′*^ values of each condition. (b) The response criteria of each condition. Values were calculated based on SDT analysis for four conditions in two groups. It indicated no significant effect on detectability and response criteria.

### fMRI results

We collected fMRI data of each participant and combined multiple approaches in the fMRI data analysis, including univariate analysis, ROI analysis, MVPA, and RSA.

#### Univariate analysis

In an exploratory analysis, to identify activated clusters associated with group differences of face conditions, we conducted a group-level analysis on the OT versus PL contrasts. We observed significant activation differences during face judgement between the OT and PL group, and the resulting statistics and coordination table was shown in Table 2. The cuneus (and occipital lobe in general) showed greater activity under all conditions for OT group, which indicated increased sensitivity to face stimuli under OT administration. Similar to previous fMRI findings (39, 51), we found stronger IFG activity in self-morphed facial conditions compared to other-morphed facial conditions, further indicating the role of IFG in self-other distinction. Moreover, the fusiform gyrus (FFG) showed stronger activity on adult-face conditions compared to child-face conditions (Table 2).

**Table 2.** Second level univariate analysis table for OT versus PL contrast.

| Conditions | Brain Region | T-score | MNI coordinates |  |  |
| --- | --- | --- | --- | --- | --- |
|  |  |  | x | y | z |
| **Self-Child** | Cuneus - L | 10.01 | -12 | -103 | 4 |
|  | Cuneus - R | 15.09 | 12 | -103 | 1 |
|  | IFG - L | 6.77 | -30 | 23 | -11 |
|  | IFG - R | 7.94 | 36 | 20 | -5 |
|  | FFG - L | 12.38 | -39 | -58 | -23 |
|  | FFG - R | 13.11 | 45 | -52 | -20 |
| **Other-Child** | Cuneus - L | 9.34 | -12 | -103 | 7 |
|  | Cuneus - R | 12.17 | 15 | -100 | 1 |
|  | IFG - L | 5.07 | -33 | 26 | 10 |
|  | IFG - R | 4.96 | 36 | 23 | -2 |
|  | FFG - L | 7.61 | -42 | -55 | -23 |
|  | FFG - R | 9.97 | 42 | -52 | -20 |
| **Self-Adult** | Cuneus - L | 10.92 | -12 | -103 | 4 |
|  | Cuneus - R | 14.35 | 12 | -103 | 4 |
|  | IFG - L | 10.41 | -33 | 20 | -2 |
|  | IFG - R | 9.80 | 33 | 23 | -2 |
|  | FFG - L | 12.22 | -42 | -55 | -23 |
|  | FFG - R | 14.04 | 42 | -52 | -20 |
| **Other-Adult** | Cuneus - L | 11.90 | -12 | -103 | 4 |
|  | Cuneus - R | 14.37 | 15 | -100 | 1 |
|  | IFG - L | 3.20 | -42 | 17 | -2 |
|  | IFG - R | 7.96 | 48 | 35 | 16 |
|  | FFG - L | 11.77 | -42 | -58 | -23 |
|  | FFG - R | 12.61 | 45 | -52 | -20 |

**Note:** Table showing regions with FDR corrected $p < 0.05$ . Coordinates were labeled using the *xjView* toolbox (https://www.alivelearn.net/xjview). IFG: inferior frontal gyrus; FFG: fusiform gyrus.

#### MVPA

We generated four MVPA classifiers to discriminate OT and PL treatments based on facial conditions (self-child/SC, other-child/OC, self-adult/SA, and other-adult/OA). For a certain classifier, if some voxels are able to significantly distinguish OT group from PL group, then it is possible that OT caused detectable neural changes over the brain regions under that condition. Fig. 5 shows voxels of each classifier that can significantly discriminate the OT vs. PL group. It revealed that voxels in the occipital cortex, MFG, and IFG of self-child/self-adult condition can classify the two groups, suggesting the roles of visual perception brain area and IFG in self-other distinction. These MVPA findings not only confirmed the univariate analysis results, but also showed that the activation pattern of more voxels in the visual cortex can classify the two groups for self-morphed child faces.

**Fig. 5.**
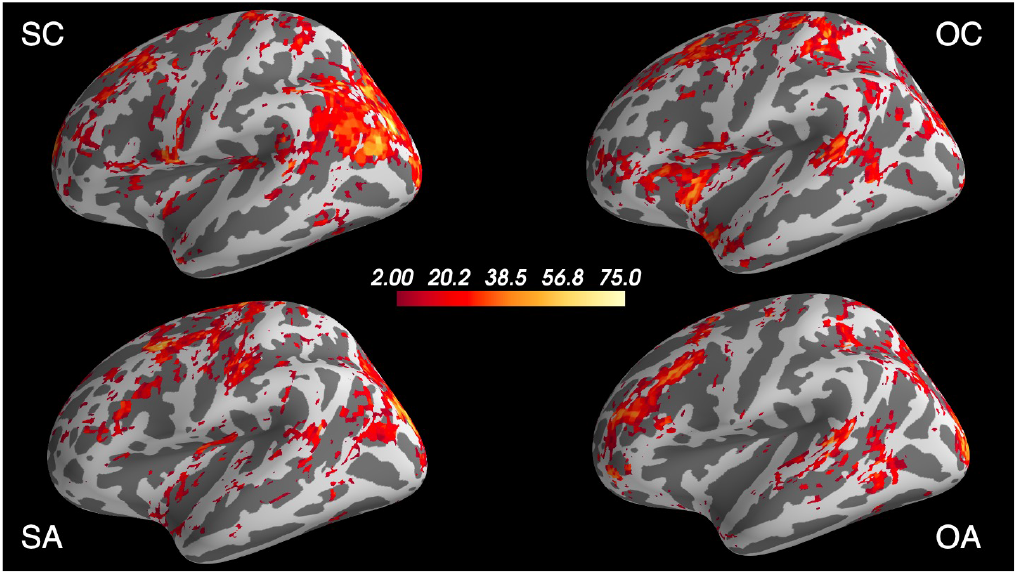
Whole brain MVPA classifier results. Values on each voxel are permuted statistics (t-value) of 500 iterations, where larger value means higher significance. A voxel is colored if the accuracy of the classifiers on discriminating OT versus PL treatment is significantly higher than the permuted baseline level. Abbreviations are consistent with previous usages. MVPA results are consistent with the univariate results, showing significance in areas such as cuneus, IFG, FFG, etc.

#### MVPA with ROIs from Neurosynth

To further examine the effect of OT on self-related processing, we tested whether fMRI data of two groups in the present study can be reliably decoded from the ROIs involved in “self-referential”, according to the Neurosynth meta-analysis. We first masked participants’ brain data with Neurosynth masks and then trained classifiers on either OT or PL group data to discriminate between self (SC and SA) and other (OC and OA) conditions. Fig. 6 (a,b) shows the significant *t*-values of the Neurosynth ROI, which statistics were generated using a similar permutation method as specified in the above MVPA analysis (Fig. 5). Despite value differences, the patterns of the significance of classifiers are similar. Therefore, a histogram of all significant t-scores in the Neurosynth ROI regions are plotted to visualize the results. As shown in the Fig. 6, the OT classifier showed more voxels with a positive t-scores, indicating that the OT classifier might be better at classifying self versus other faces. This indication further implied that OT treatment might increase the activation difference between self and other conditions, resulting in the better performance of the OT classifier.

**Fig. 6.**
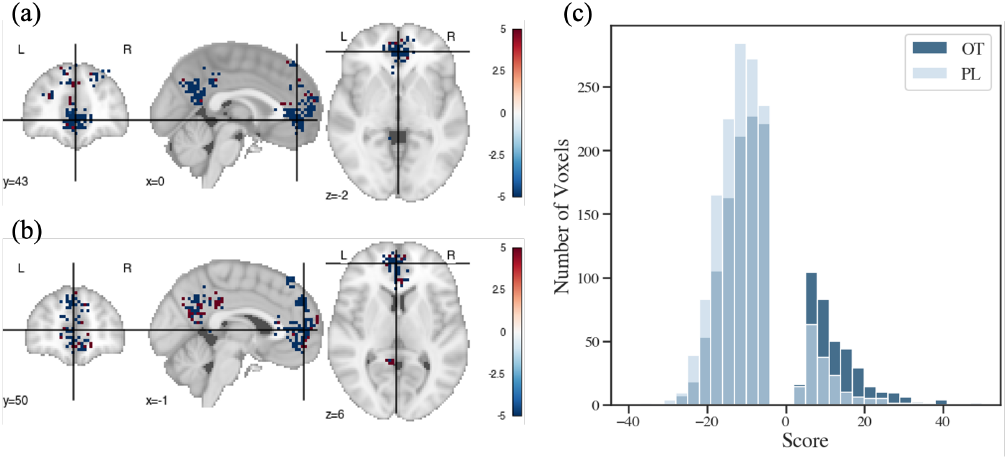
MVPA results with masks from Neurosynth self-referential ROI. (a) Statistic value of OT classifier over the self-referential ROI. The values of statistics scores shown on the plot were capped at ± 5 for better visual representation. (b) Statistic value of PL classifier over the self-referential ROI. (c) Histogram of statistic scores in Neurosynth masked MVPA results. Only significant statistics were counted, causing the frequency around zero to be 0.

### ROI results from AAL

To compare the activation under each condition, we extracted the average PSC of each ROI of each condition for further statistical analysis. Sixteen ROIs, which have been shown contribute to self-other discrimination (coordinates from (52)) and facial recognition (coordinates from (43)) were defined using the AAL3 template. The locations of the ROIs selected were shown in Fig. 7. PSC values were calculated under each condition to identify the difference between self and other faces for two groups. For plots of other selected ROIs and detailed statistic information, see supplementary material Fig. S2, S3, S4. Fig. 8 shows the PSC plots of the bilateral ACC, IFG-operant part, IFG-triangular part, and insula. It showed that OT increased self-other differentiation, particularly for adult faces, which effect was more pronounced in left hemisphere regions. With self/other face and child/adult face as two factors, the two-way ANOVA on PSC of IFG revealed significant influence from self/other face (OT: *p <* 0.0001, PL: *p* = 0.0005) but not on child/adult face (OT: *p* = 0.2315, PL: *p* = 0.1297). Other ROIs showed the similar pattern, especially over the left hemisphere.

**Fig. 7.**
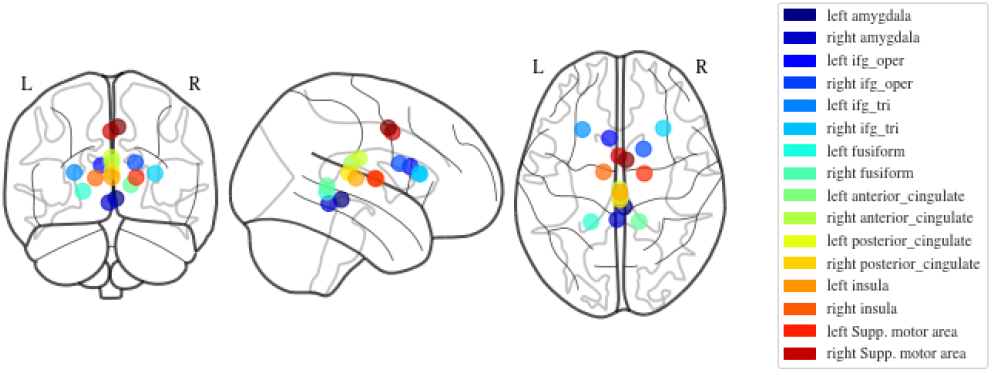
Visualization of center of mass locations for each selected ROIs from the AAL3 template. Note that this is a rough representation of the location of the ROIs, calculated by averaging coordinates in each of the ROIs.

**Fig. 8.**
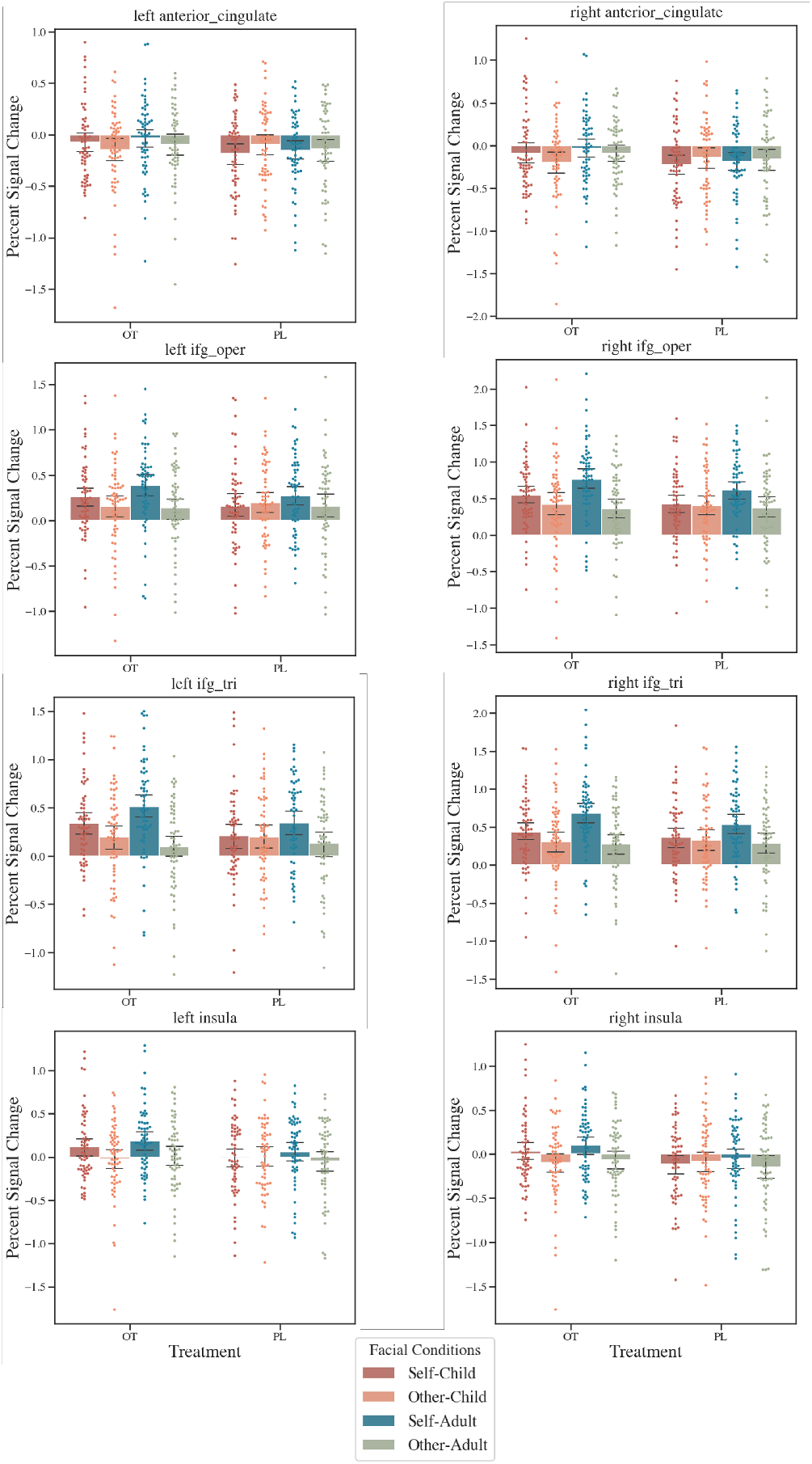
Percent Signal Change over IFG (in AAL3 template) of the ROI analyzed. ROI names are noted in the titles of the sub-figures. Error bars show the boot-strapped confidence intervals with 500 iterations.

### Representational connectivity difference between OT and PL group

To explore the possible correlations between activation of the prior ROIs, we calculated the similarity of the activation for each ROI in both OT and PL groups respectively. With Fisher-Z transformation, statistics for correlation difference between RDM correlation of OT and PL were calculated. The significance of OT PL difference is shown in Fig. 9. As shown in the figure, there was ROI connectivity difference (e.g., amygdala and ACC, FFG and insula) between groups. The result suggests a higher representational connectivity over OT administration group in these face and self-referential ROIs.

**Fig. 9.**
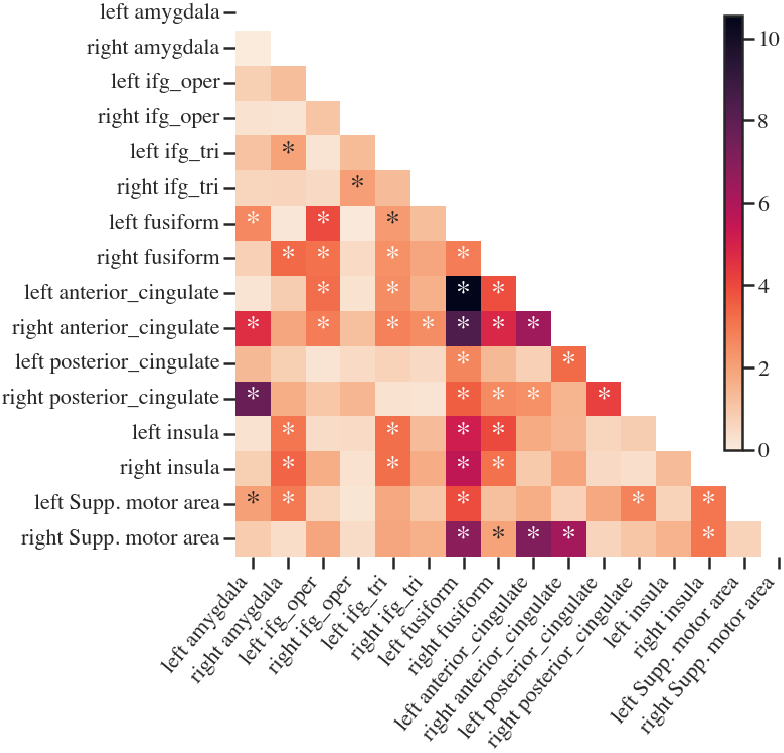
Z-scores of connectivity-similarity difference between OT and PL group. Statistics for correlation difference between RDM correlations of OT and PL, using Fisher-Z transformation. Labels on the axis are the names of the selected ROIs, the same as they appeared in the AAL3 atlas. An asterisk (“*”) indicates significance in that pair of correlation difference.

### Correlations among behavioral, psychometric data and fMRI data

Up to this point, we have displayed the results of three types of data: psychometric data, behavioral data, and fMRI data. We were able to derive different implications from these data of different modalities. In this section, we would try to bridge the data and provide interpretations of their relationships.

#### Correlation between ROI activation

Fig. 10 shows the difference of OT groups’ and PL groups’ correlation among ROI activation. The ROI-ROI correlations indicate the coactivation pattern in the self and face perception brain regions. The figure showed that more pairs of ROIs were significant in the adult face conditions (“self-adult” and “other-adult”). This indicates that compared to child-morphed stimuli, OT might modulate adult-morphed faces, and result in changing co-activation pattern in face related and self related ROIs.

**Fig. 10.**
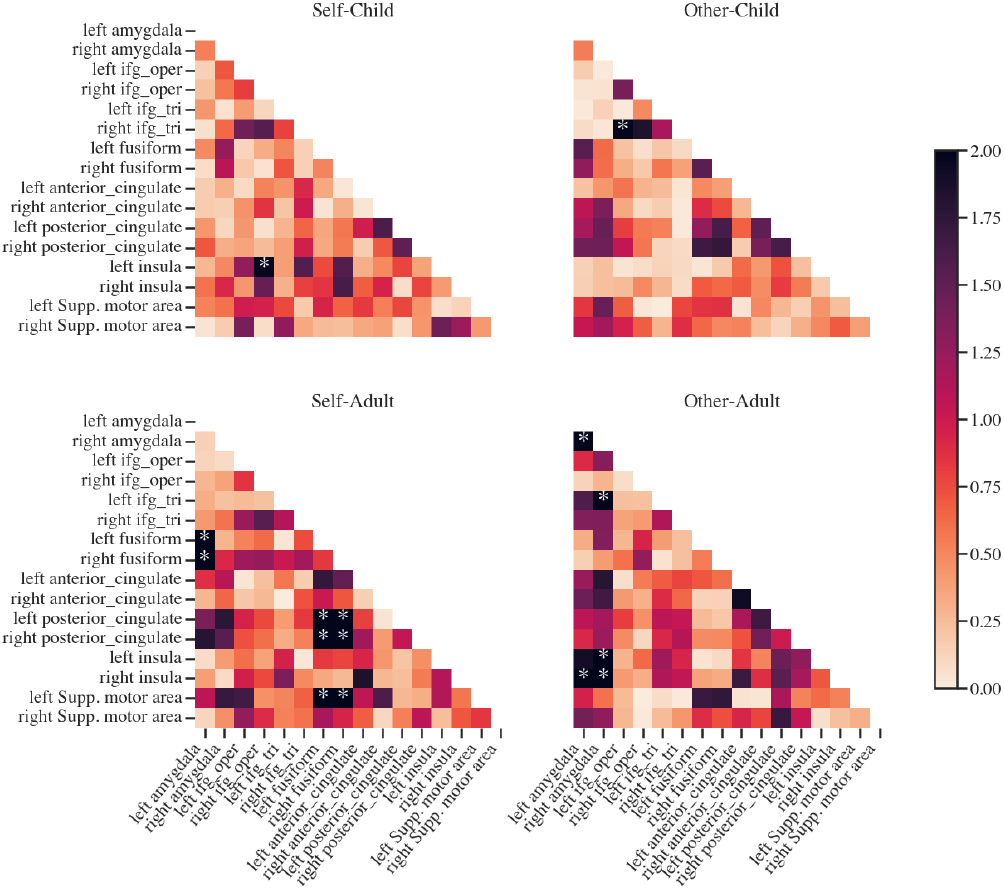
Significance of correlation differences between OT and PL group, divided by the four facial conditions. Significance were calculated using the Fisher-Z transformation. Asterisk (∗) indicates significance.

#### Correlation between psychometric data and fMRI data

Since we have found interesting OT effect on self-other differentiation based on fMRI data, to investigate the relationship between brain activities and psychometric scores in participants, we conducted bivariate correlation analysis between behavioral scores and PSC activity difference (self-other) over selected ROIs (similar as difference calculation in the ROIROI analysis). We first calculated the correlations between behavioral data and neural activation of self versus other condition respectively. Based on experimental design and previous analysis on psychometric data, OT and PL group difference is not considered. Then, similar to previous process, we derived the difference between self and other and calculated the significance. Fig. 11 shows the z-scores of the difference. The only significant difference appeared in child condition: the correlation difference between left insula and Machiavellianism Scale (from psychometric data) was significant. This is consistent with previous researches, where individuals with different Machiavellianism Scales scores exhibit different volume and activation in the insula areas (53, 54).

**Fig. 11.**
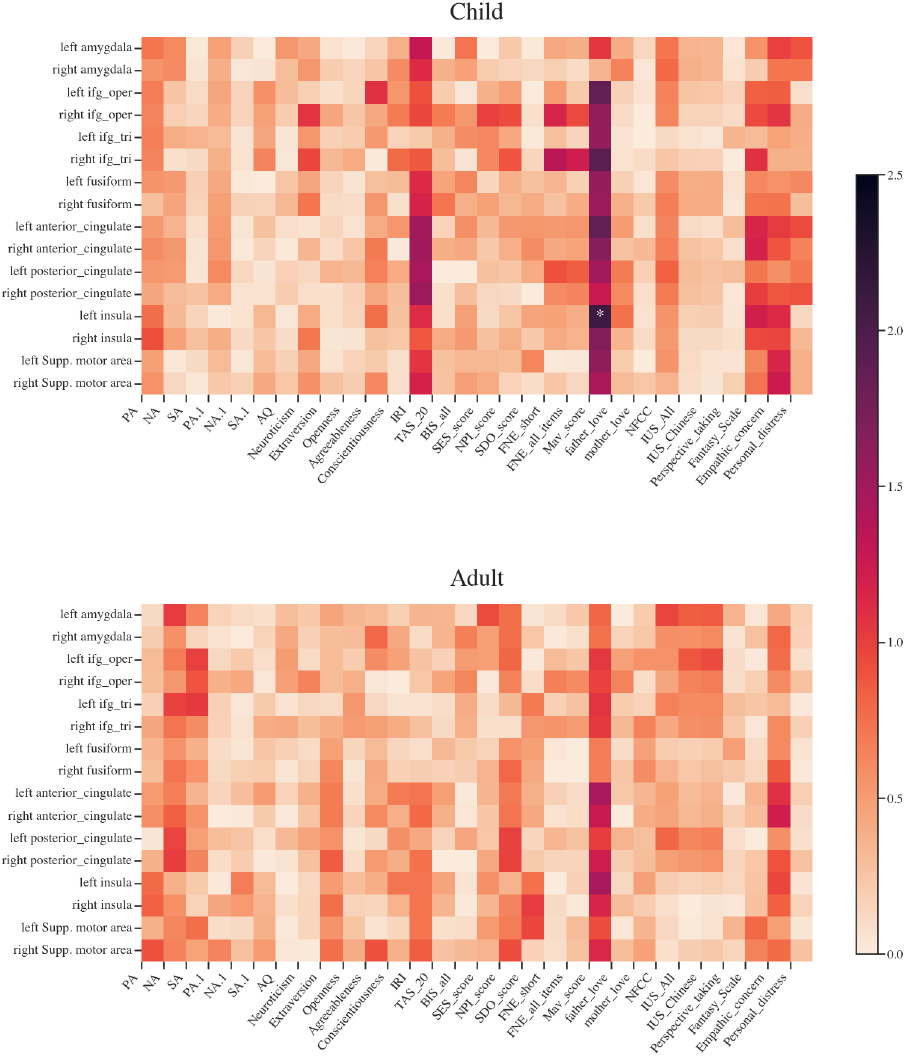
Significance of correlation differences between self condition and other condition, divided by child face and adult face condition. Asterisk (∗) indicates significance.

Although administering different treatments have little influence on psychometric data, we have shown that treatment can have influence on fMRI activations. Therefore, the correlation patterns of psychometric data and ROI activation between OT and PL treatment could be different. As shown in Fig. 12, the difference of correlation between OT and PL group was more significant in other-face conditions (other-child and other-adult) compared to self-face conditions. Specifically, the IRI showed more significant difference in other-face conditions. We found that in OT participants, IRI have higher correlations with the ROI activations (see Fig. 12). This trend showed not only in other-face conditions, but also in self-face conditions, albeit less significant. A possible explanation is that OT modifies the activation in the selected brain areas, and consequently their correlation with the IRI scores. It is possible that brain regions of participants under administration of OT would activate more differently when seeing a other-morphed face, resulting in a more significant difference.

**Fig. 12.**
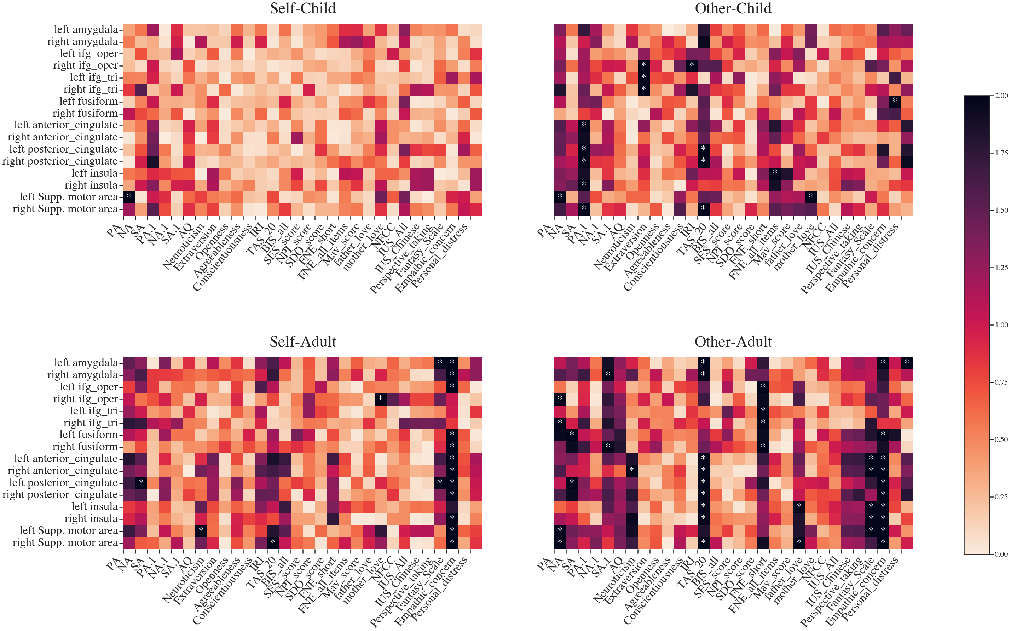
Significance of correlation differences between OT and PL group. Asterisk (∗) indicates significance. Separate correlation heatmap of OT and PL group are shown in Fig. S7.

#### Principal component analysis (PCA) on the whole brain activation

Except from representing the fMRI by voxel and by ROI, PCA could also be used in brain imaging data to derive a principal pattern of the activation. As we previously mentioned, principle component of the brain activations could easily be calculated with linear algebra in the python sklearn package (47). The first principle component was able to explain 23.42% of the variance among all the participants. The upper left corner of Fig. 13 shows the principle component on the whole brain. By projecting each participant’s activation pattern onto the first principle component, we derived a projection score for each participant. We used the Boruta package in R (55) to calculate the contribution of each questionnaire score to the projection (Fig. 13). Among all of the variables, scores regarding intolerance of uncertainty (“IUS_all” and “IUS_Chinese”) and social anxiety showed significant contribution. The result indicates that the found principle component might be explained with uncertainty and anxiety traits in individuals.

**Fig. 13.**
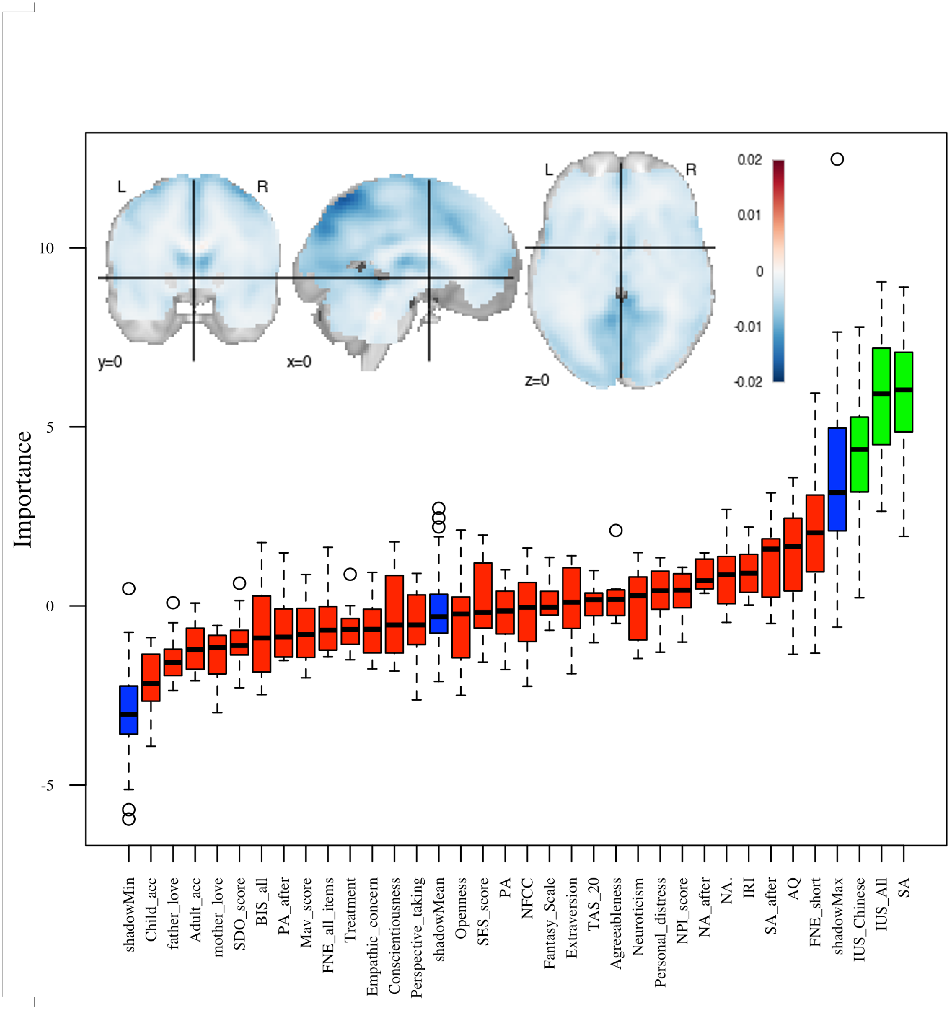
Variable importance plot, showing psychometric variables’ contribution to the first principle component of fMRI data. Brain pattern in the upper right corner shows the values of each voxel in the first principle component. Green boxes indicate significant importance, and red boxes indicates non-significant importance. Asterisk (∗) indicates significance.

## Discussion

Accumulative evidence indicated that the OT influences human behavior (56) and brain activities (57–59). Since the previous behavioral results are mixed on whether OT sharpens or blurs self-other distinction (36, 60, 61), in the present study, participants make judgements in the self-other differentiation task with morphed faces. To our knowledge, this is the first fMRI study to investigate the effect of OT on self-other differentiation with self-morphed adult and child faces, with comprehensive behavioral and neural analysis.

Although we did not observe significant OT effect on behavioral responses, we were able to show that: (1) OT increases brain activities in self- and face-related brain regions, specifically in the IFG and visual areas, when they perceive faces, (2) the voxels over visual cortex for self-morphed child faces can classify OT vs. PL group, (3) the OT vs PL self-other differentiation effect in brain activity is more pronounced for adult faces in the left hemisphere.

### OT effects on self-relevant face processing in neural level but not on behavioral performance

The psychometric data were collected to control and observe the homogeneity of the two groups (OT and PL). As a result, we predicted that there would be no significant difference on those variables between the two groups, except for the potential significance of those variables that were tested before and after the treatment (such as PA and PA_after). Our results on psychometric data did fit our initial prediction on it.

For behavioral performance, previous study indicated that OT reduced RTs for making both self and other judgments (36), and reduced the recall accuracy (37). We did not observe significant OT modulation effect on accuracy(Fig. 2a), RTs (Fig. 2b) and detectabliliy in SDT analysis(Fig. 3) in the current self-other differentiation task. Even though non-significant, our behavioral analysis showed that the OT group showed better diagnostic ability than the PL group (Fig. 3). It is possible that OT may facilitate overall behavioral performance in the current task. Moreover, in the behavioral results, we also observed significantly better performance for adult-morphed facial stimuli (Fig. 2). This might be attributed to the dominant age effect that the better performance toward adult facial stimuli, along with the generally higher accuracy and shorter responses time.

However, for our fMRI results, we observed general increased brain activation over visual area (cuneus) and IFGs(Table 2) for OT group compared to PL group. As proposed by previous literature, facial features are first encoded by visual area, and after this process, the frontal cortex encoded self-referential properties (62). Therefore, our second-level result indicating brain regions in processing of faces within the occipital cortex(fusisorm gyrus and cuneus) is consistent with the work of processing of faces with regions within the fusiform gyrus (63–65). Based on Uddin et al. (2005)’s report that the IFG in the right hemisphere responded parametrically to the amount that a facial stimulus looked like the participant (66), the increased IFG activity for OT group in our study suggest more self-resemblance processing after OT administration. The ROI analysis and the MVPA results further confirmed that there were significant difference in brain regions such as the IFG. For the ROIs in face processing and self-referentail processing, we further observed greater self-other distinction in ROI activity, especially in adult-morphed faces. For example, we find IFG and insula activities are greater for self-adult than other-adult. Inferior parietal cortex along with the prefrontal cortex comprise a self–other brain network, which is important in distinguishing the self from the other, and the shared self-other discrimination is considered as root for prosociality such as empathy and trust (67). Interestingly, previous works showed that viewing self face activate the IFG, inferior parietal lobe and inferior occipital cortex, especially in the right hemisphere (66, 68, 69). However, in our study, we found that the OT group showed stronger self-other distinction in the left hemisphere. Suppose there is a right hemisphere advantage in “self” network, we provide first evidence of OT sharpening the left hemisphere self network, supporting greater self-other distinction. Thus, our ROI results showing higher self-other distinction might reflect pronounced role of left hemisphere area, which may be responsible for maintaining self–other distinctions under OT administration. It may also reflect a flexibly adaptation to update self-other distinction after OT administration.

Consistent with this possibility, the MVPA results provide further evidence for our explanation. It first confirmed that the face processing and self-referential processing brain regions can classify two treatment groups. Furthermore, the results indicate the OT can enhance classification between self-morphed and other faces, in the voxels of self-referential ROIs. Given that OT increased self-other distinction in the left hemisphere, the better classification performance in self ROIs seems more plausible. Supporting the higher social salience or flexibility in OT group, it demonstrate further evidence relevant for social cognition that such flexibility may be due to hemisphere balances.

### Child face vs. adult face: higher OT effect on adult faces

It is notable that the self-other distinction was stronger in adult-morphed faces rather than child-morphed faces, regardless of the treatment condition (OT and PL) (Fig. 2a, Fig. 3). This overall adult-face advantage could reflect an own-age bias (1), such that the recognition for faces of one’s own age group is often better than that for faces of another age group(70). Accordingly, with respect to the the age cohort of participants (adults), the higher performance of adult faces across two groups reflect the typical advantage in self-age face processing. Researches have also shown that compared to outgroup faces, ingroup faces receive more holistic and in-depth processing, which facilitates the perceptual discrimination of faces (71, 72). Participants of the current study were all adults, hence showing an ingroup tendency toward adult faces. Our results dovetail nicely with the above findings.

Another possible account of this results is that people have various capacity in recognizing one’s own current and past facial appearance (73). Studies suggest different brain regions in processing one’s current and childhood face (73). The inferior occipital gyrus, the superior parietal lobule and the inferior temporal gyrus are more involved in current self-processing, while TPJ and the inferior parietal lobule, are more involved in childhood self-processing. According to this account, OT may only affect the self-other differentiation in recognition of the current self-face, but not past self-face, as the latter requires more memory encoding and retrieval (74).

Therefore, the better detection accuracy in the adult face condition is consistent with our previous work (1), and may reflect the automatic processing of self-age face. For adult participants, a child-morphed stimulus may require for the retrieval and maintenance of childhood self images, as the image is not their current facial appearance. Thus, there is a more indirect link between the childhood image and the response “self” than adult-morphed faces and “self”. It would lead to the lower accuracy, longer RTs and no OT enhancement effect in the brain activity. In this vein, how OT would modulate the self-face recognition across ages is still an open question, but the current different effects on self-other distinction in adult and child face may yield important insights into how OT modulates social processing of faces with different ages.

### Individual differences of the OT effect: Brain-behavior association

Following previous work showing individual difference on the OT effect, our findings also offer novel associations between brain activity, functional connectivity and personality traits. A recent work has suggested the OT effect on self-other distinction is influenced by oxytocin receptor (OXTR) genotype (61), as the OT only modulates the task decision time of rs53576 G carriers. Although we do not have the gene sequencing data of participants, it is possible that similar link between individual difference and the effect of OT. In the analysis on psychometric data, we found correlations between participants’ psychometric questionnaires scores and functional connectivities in the fMRI task, which indicate the effect of OT on self-brain might be modulated by individual difference.

For example, at correlation level, we observed significant brain-behavior correlation (IFG activity and measured IRI score) difference. It means that there is higher IRI score and IFG activity correlation for the other-child condition of OT group than PL group. Since self-other overlap is critical for empathy, it echos studies that showed correlation between personal empathy trait and self-other overlap (75–77). OT, to some extent, strengthen this link of individual difference for social adaptation (78). That is, after OT administration, people with higher IRI might enhance self-other overlap, while people with lower IRI might enhance self-other distinction. Together with other correlations, these findings support the view that differential pharmacological effects have roots in individual differences in personality traits.

### Limitations and future perspective

There are several limitations that should be addressed. First, current study is a between-subject design, which is unable to track the neural response change before and after OT administration directly. Second, the sample size of the participants and the male-only participant group may not allow generalization of the conclusion. Since there are gender differences in the effects of stress and OT on self-other distinction (79), future research could utilize bigger sample with more diverse participants and probably capture gender difference in the effect of OT. It could combine with pre- and post-administration tests to investigate the large-scale neural-network, and to track and reveal the underlying neural mechanism of OT effect in self-other distinction processing.

The present findings substantially extend previous findings on the effect of OT on neural response to social salient stimuli (80). Our study first used self-morphed faces to investigate the OT effect on self-relevant processing, more specifically tap into self-other distinction. Evidence have shown distinct neural underpinnings of the processes of self- and other-related information, which is critical for human social motivation and behaviors(81). A previous study using trait judgement task showed OT effects on self-referential processing, including reduced RTs for self-related trait judgments, increased accuracy in memory retrieval, and decreased MPFC activation in self-related trait adjectives (82). Although our findings show no significant OT effects on the behavioral judgments and RTs, the comprehensive fMRI analysis results indicated the robustness of increased self-other distinction in OT group, especially for adult faces.

## Conclusions

Taken together, the current findings extend previous research linking OT effect and self-relevant information processing, which provide potential core mechanisms associated with OT modulation effects on social behaviors. Our results highlight significant larger self-other difference in brain activities, particularly within the left face and self -referential processing brain regions. This suggests a possible OT induced complementary left self-brain network. These OT effects were associated with personality traits which confirm the individual difference in OT effects of self-other distinction brain activity. It may shed lights on the roles of self-relevant processing on social behavioral change under OT administration.

## Ethical Statement

The authors are accountable for all aspects of the work in ensuring that questions related to the accuracy or integrity of any part of the work are appropriately investigated and resolved. All procedures performed in this study involving human participants were in accordance with the Declaration of Helsinki (as revised in 2013). The ethics committee of Beijing Normal University approved this study and written informed consents were obtained from all participants.

## Data and Code availability

The data is available upon request to the corresponding author. The code for data analysis is available at https://github.com/andlab-um/OT_face.

## Acknowledgements

This work was mainly supported by the Science and Technology Development Fund(FDCT) of Macau [0127/2020/A3], SRG of UM(SRG2020-00027-ICI), in part by the Natural Science Foundation of Guangdong Province(2021A1515012509). The authors would like to thank Qingyuan Wu, Shensheng Wang, Yanghua Ye, Quanying Liu and Kun Chen for comments on the early version of the manuscript. We also thank all research assistants who provided general support in participants recruiting, and data collection.

## Author Contributions

H. W. designed research; H. W. performed research; Y. W. and H.W.analyzed data; and Y. W., R. W., and H. W. wrote the paper.

