## Supplementary for "The role of oxytocin in modulating self-other distinction in human brain: a pharmacological fMRI study"

**This PDF file includes:**

Figs. S1 to S7
SI References

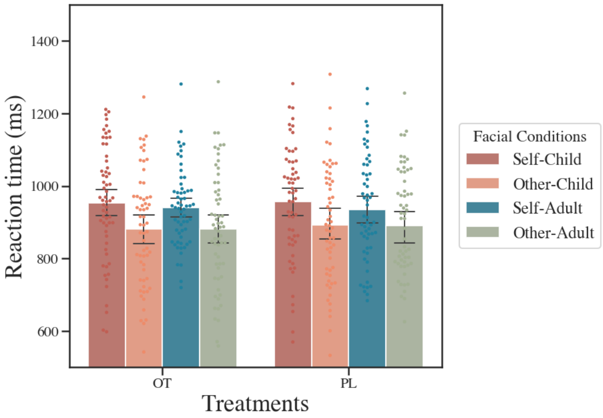


**(a)** Reaction time, facial condition decided by actual stimuli.

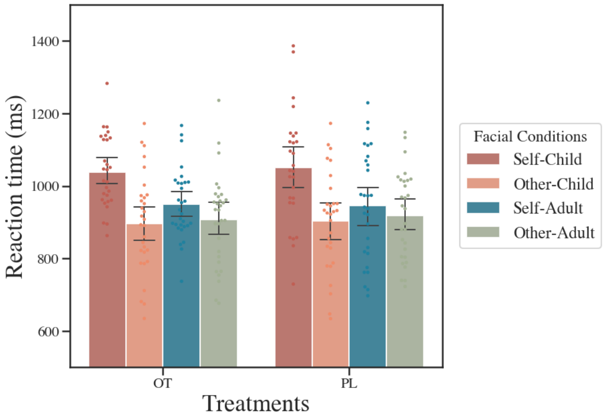


**(b)** Reaction time, facial condition decided by participants' response.

**Fig. S1.** Reaction time of participants, separated by facial conditions. (a) The plot that is used in main manuscript. Facial conditions are determined by the actual stimuli. (b) Facial conditions determined by participants' response.

During analysis of reaction time, we realized that participants' subjective perception (i.e. whether they perceive the stimuli as themselves, regardless of the actual origin of stimuli) could also influence their judgement. Therefore, we performed a secondary reaction time analysis, where the the facial conditions are determined by whether participants reported as self-face in the discrimination task. If that is the case, then we assign "self", accompanied by the either "child" and "adult", depending on what the face was actually morphed with, as the facial condition of this stimulus. Similar process of facial condition assignment is applied for the stimuli that participants reported as other-face. We observed that the differences of reaction time for child faces increased in both OT and PL treatment group. Adult-morphed faces, on the other hand, did not show a similar trend.

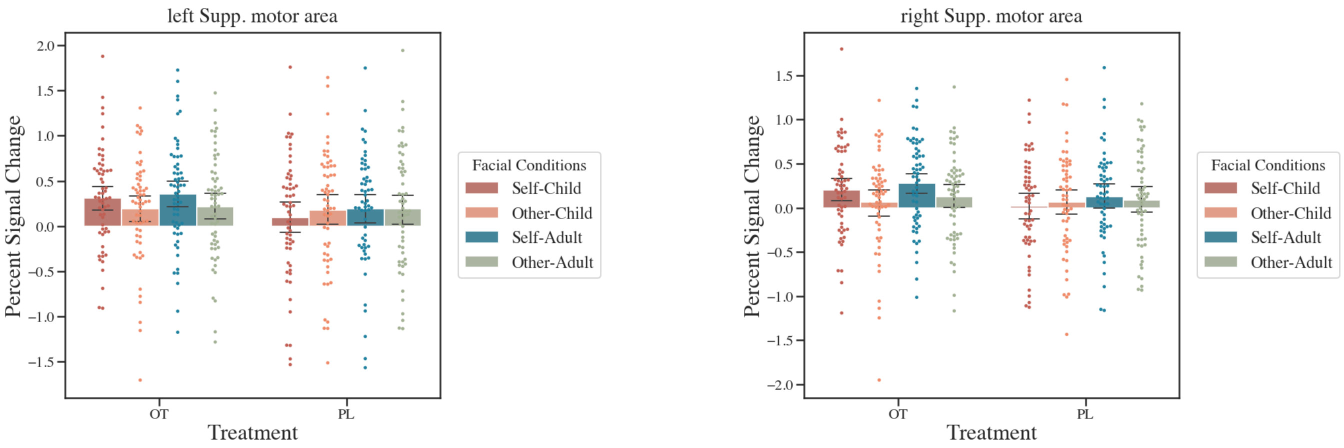


**Fig. S2.** In the ROI analysis of fMRI data, we also included the supplementary motor area. The initial intention was to pick a "neutral" region and see if similar trend as in previous ROI analysis would occur. Interestingly, in both supplementary motor area in left and right hemisphere, the influence of self-face was significant in the OT group, and insignificant in PL group (Fig. ??). This change in significance might indicate that OT is related to activation pattern differences in supplementary motor area. Previous literature mentioned that interpersonal motor coordination (IMC) might be associated with interpersonal behaviors (1). Previous studies of neural imaging evidence regarding IMC focused on the neural coupling and correlation between brains and brain activation (2, 3). It is possible that supplementary motor area is related to the process of IMC: OT amplifies the potential of IMC happening based on the given morphed face, and the activation of supplementary motor area is also modified.

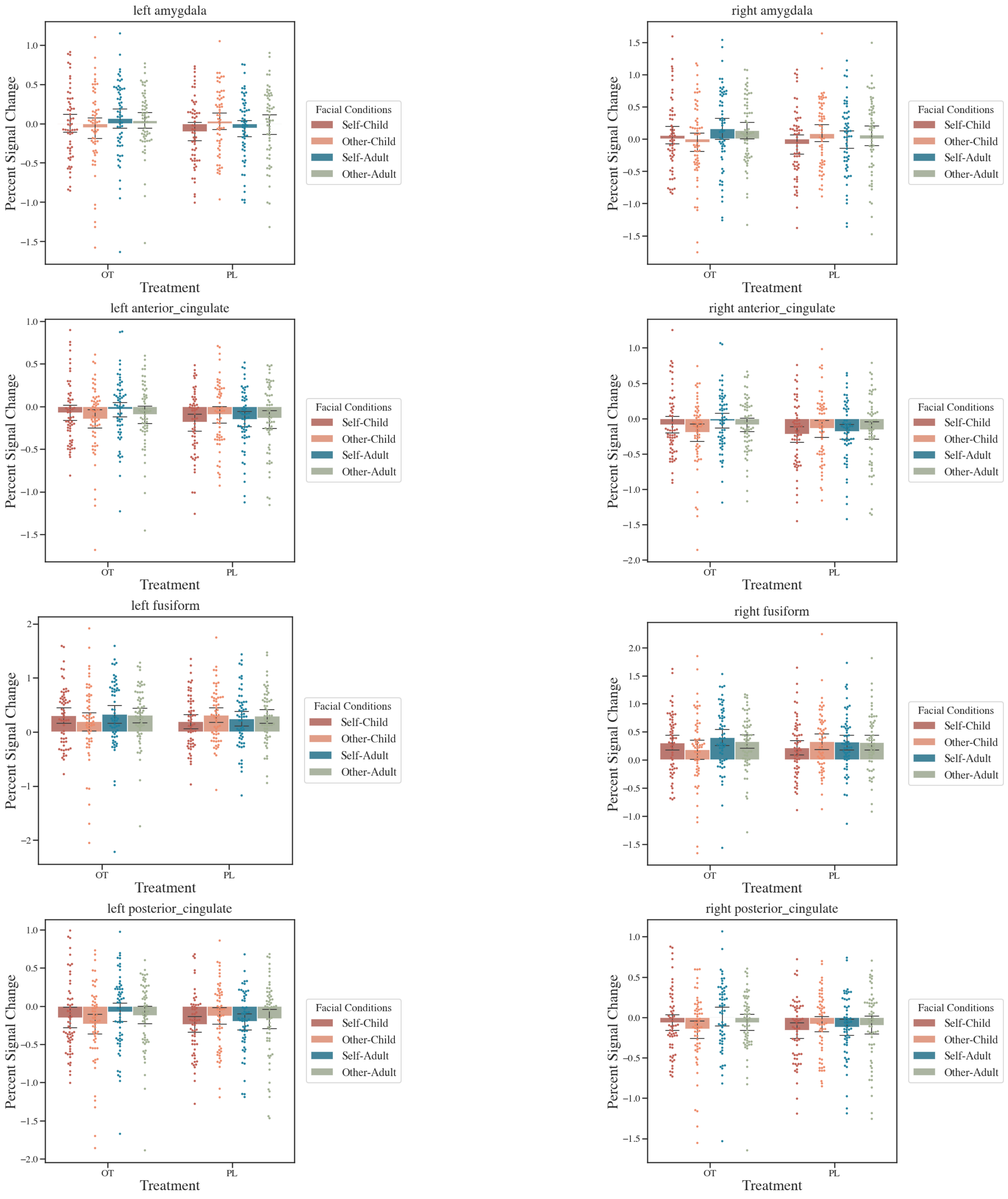


**Fig. S3.** All other PSC from considered regions. Similarly, regions are indicated by the titles of subfigures.

| ROI | Treatment | Child face: p | Self face: p | Interaction: p | sa–oa difference: t–test |
|---|---|---|---|---|---|
| left amygdala | OT | 0.2784 | 0.376 | 0.7631 | 0.2735 |
| left amygdala | PL | 0.9913 | 0.011 | 0.294 | |
| right amygdala | OT | 0.0801 | 0.2106 | 0.4237 | 0.3266 |
| right amygdala | PL | 0.6373 | 0.0063 | 0.2511 | |
| left ifg_oper | OT | 0.3569 | 0.0003 | 0.0469 | 0.0834 |
| left ifg_oper | PL | 0.4161 | 0.2836 | 0.1209 | |
| right ifg_oper | OT | 0.2315 | 0 | 0.0009 | 0.0459 |
| right ifg_oper | PL | 0.1297 | 0.0005 | 0.0397 | |
| left ifg_tri | OT | 0.5882 | 0 | 0.0006 | 0.0172 |
| left ifg_tri | PL | 0.5022 | 0.0024 | 0.0338 | |
| right ifg_tri | OT | 0.1171 | 0 | 0.0013 | 0.0675 |
| right ifg_tri | PL | 0.2157 | 0.0016 | 0.0221 | |
| left fusiform | OT | 0.356 | 0.2306 | 0.3393 | 0.4331 |
| left fusiform | PL | 0.7241 | 0.0318 | 0.4166 | |
| right fusiform | OT | 0.0913 | 0.0596 | 0.6236 | 0.3575 |
| right fusiform | PL | 0.423 | 0.173 | 0.176 | |
| left anterior_cingulate | OT | 0.4529 | 0.0846 | 0.8496 | 0.2501 |
| left anterior_cingulate | PL | 0.9613 | 0.1093 | 0.2765 | |
| right anterior_cingulate | OT | 0.2462 | 0.0718 | 0.5475 | 0.2796 |
| right anterior_cingulate | PL | 0.845 | 0.1352 | 0.4447 | |
| left posterior_cingulate | OT | 0.2277 | 0.1404 | 0.6614 | 0.2937 |
| left posterior_cingulate | PL | 0.9554 | 0.0374 | 0.3663 | |
| right posterior_cingulate | OT | 0.2628 | 0.0703 | 0.9328 | 0.1626 |
| right posterior_cingulate | PL | 0.8418 | 0.1223 | 0.4219 | |
| left insula | OT | 0.3989 | 0.0021 | 0.5153 | 0.3402 |
| left insula | PL | 0.893 | 0.184 | 0.1539 | |
| right insula | OT | 0.413 | 0.003 | 0.5385 | 0.3956 |
| right insula | PL | 0.9715 | 0.3403 | 0.1125 | |
| left Supp. motor area | OT | 0.6784 | 0.0161 | 0.8617 | 0.163 |
| left Supp. motor area | PL | 0.4767 | 0.4496 | 0.4803 | |
| right Supp. motor area | OT | 0.3672 | 0.0023 | 0.8085 | 0.166 |
| right Supp. motor area | PL | 0.3735 | 0.9013 | 0.3676 | |

**Fig. S4.** Statistics of PSC comparisons in all selected ROIs. The last column is the p-value for sa-oa differences. The three columns in the middle ("Child face: p", "Self face: p", and "Interaction: p") indicates the p-value for each factor from ANOVA analysis on the ROI. The sa-oa (SelfAdult-OtherAdult) difference was calculated in both OT and PL group. The a t-test was performed for each of the ROI, and the p-value (without correction) is reported in the last column.

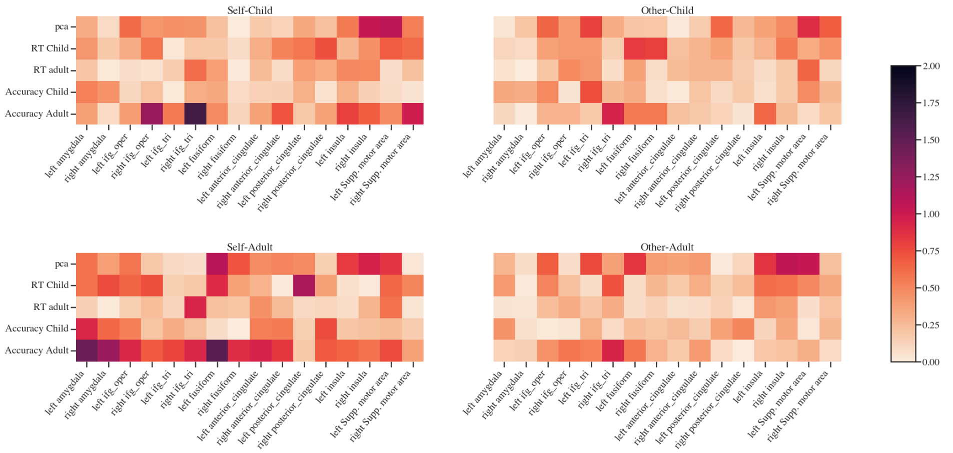


**(a)** Correlation difference between OT and PL group of behavioral data and ROI activation

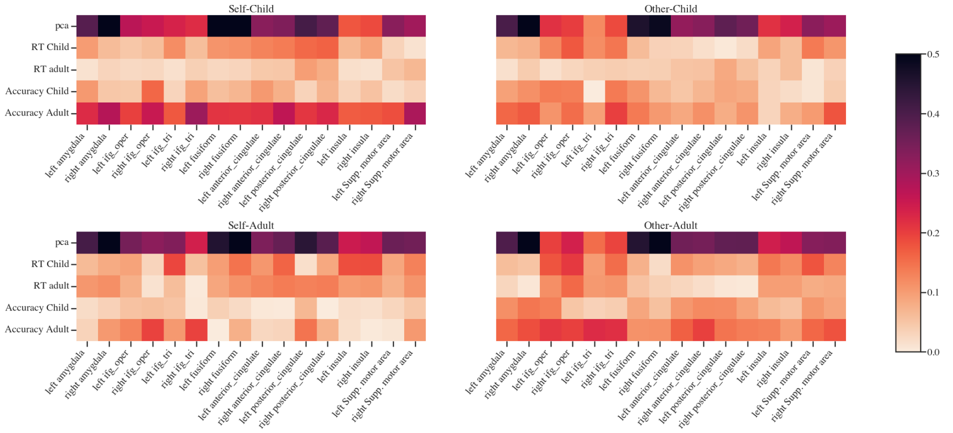


**(b)** Correlation of behavioral data and ROI activation, OT group

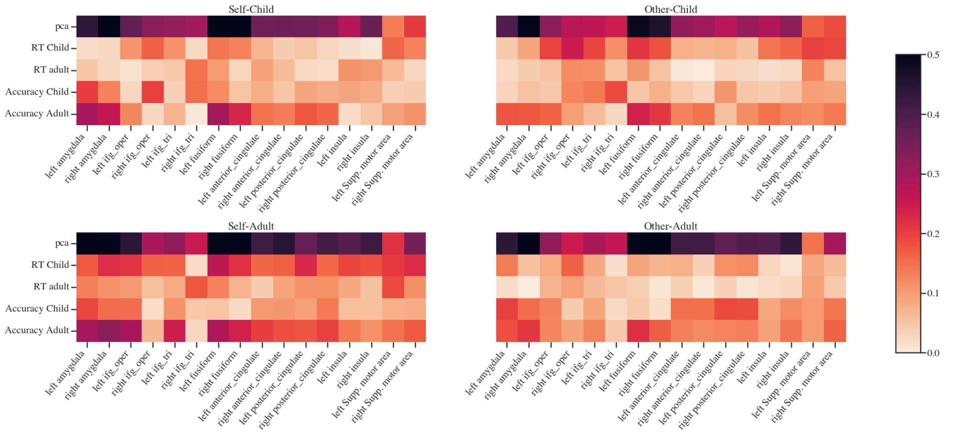


**(c)** Correlation of behavioral data and ROI activation, PL group

**Fig. S5.** Correlation between behavioral data and ROI activation. Even though not a behavioral measure, "pca" indicates the brain's projection onto the first principle component. None of the differences between OT and PL group is significant. Note that though with different patterns, both OT and PL group have low correlation. This might contribute to the insignificance from the difference analysis.

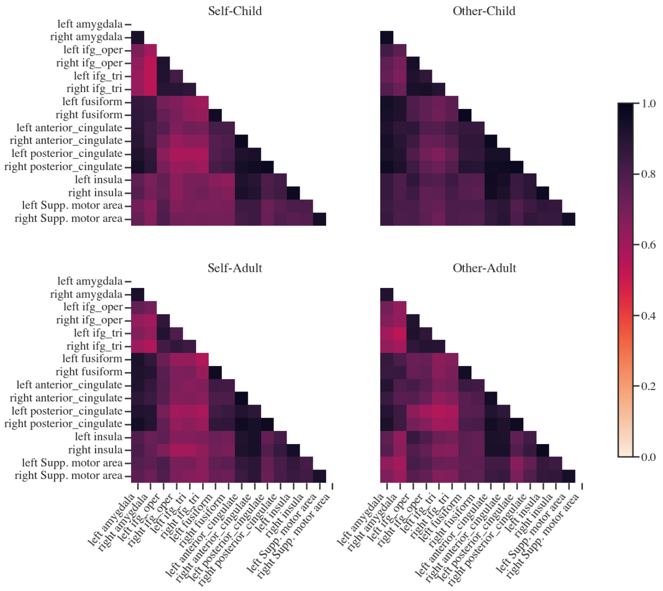


**(a)** Correlation matrix of selected ROIs, OT group

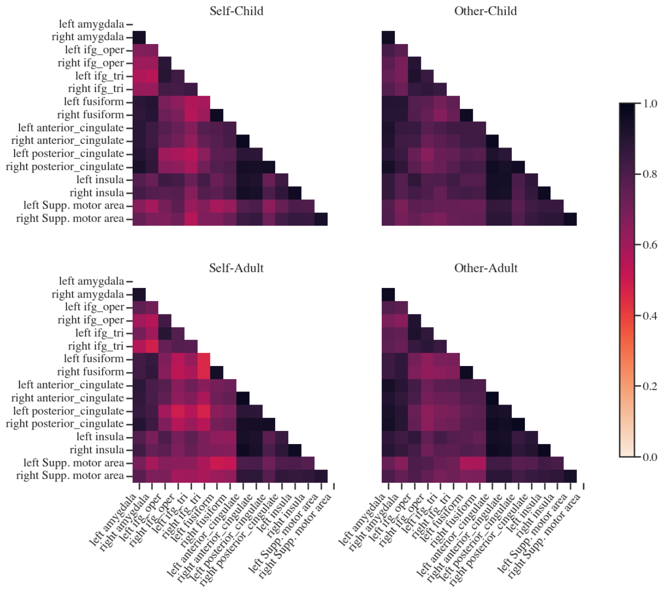


**(b)** Correlation matrix of selected ROIs, PL group

**Fig. S6.** Separated correlation heatmap of ROI activation for OT and PL group. All the ROI pair have relatively high correlation, in both OT and PL treatment groups. Therefore, a difference significance plot as shown in the main manuscript is a clearer representation of this analysis.
Despite the similarities, we could observe different patterns in OT and PL group.

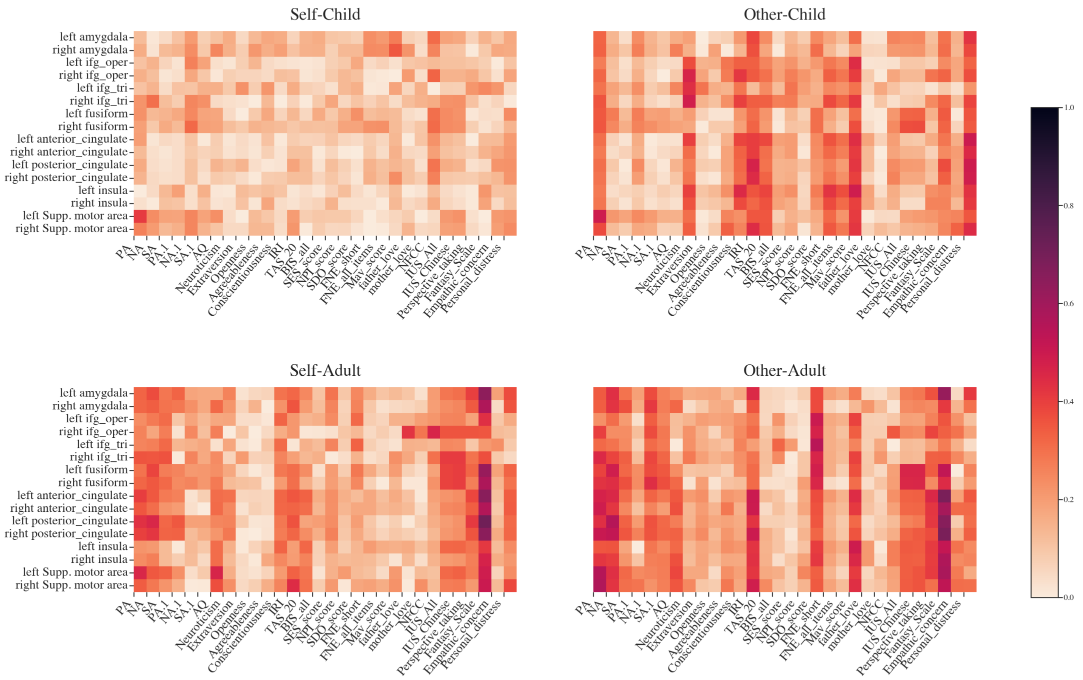


**(a)** Correlation of psychometric data and ROI activation, OT group

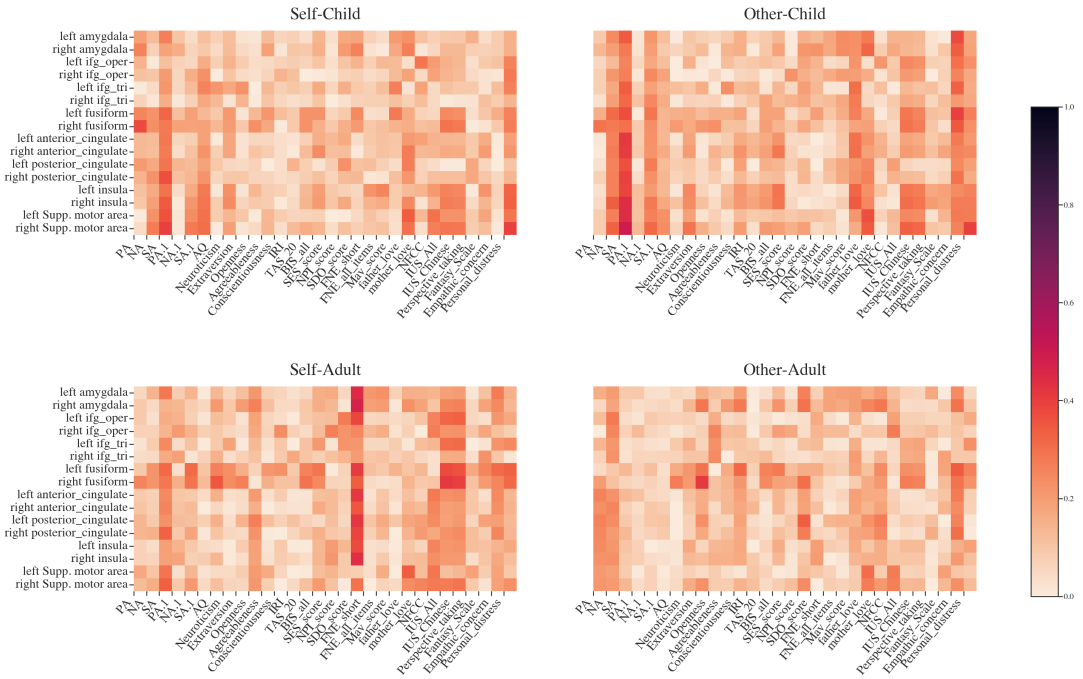


**(b)** Correlation of psychometric data and ROI activation, PL group

**Fig. S7.** Separated correlation heatmap of psychometric data and ROI activation for OT and PL group. In general, OT group have higher correlations than PL group.
